# Prediction of anti-freezing proteins from their evolutionary profile

**DOI:** 10.1101/2024.04.28.591577

**Authors:** Nishant Kumar, Shubham Choudhury, Nisha Bajiya, Sumeet Patiyal, Gajendra P. S. Raghava

## Abstract

1.

Prediction of antifreeze proteins (AFPs) holds significant importance due to their diverse applications in healthcare. An inherent limitation of current AFP prediction methods is their reliance on unreviewed proteins for evaluation. This study evaluates proposed and existing methods on an independent dataset containing 81 AFPs and 73 non-AFPs obtained from Uniport, which have been already reviewed by experts. Initially, we constructed machine learning models for AFP prediction using selected composition-based protein features and achieved a peak AUC of 0.90 with an MCC of 0.69 on the independent dataset. Subsequently, we observed a notable enhancement in model performance, with the AUC increasing from 0.90 to 0.93 upon incorporating evolutionary information instead of relying solely on the primary sequence of proteins. Furthermore, we explored hybrid models integrating our machine learning approaches with BLAST-based similarity and motif-based methods. However, the performance of these hybrid models either matched or was inferior to that of our best machine-learning model. Our best model based on evolutionary information outperforms all existing methods on independent/validation dataset. To facilitate users, a user-friendly web server with a standalone package named “AFPropred” was developed (https://webs.iiitd.edu.in/raghava/afpropred).

**Highlights:**

- Prediction of antifreeze proteins with high precision
- Evaluation of prediction models on an independent dataset
- Machine learning based models using sequence composition
- Evolutionary information based prediction models
- A webserver for predicting, scanning, and designing AFPs.

**Author’s Biography:**

1. Nishant Kumar is currently working as Ph.D. in Computational biology from Department of Computational Biology, Indraprastha Institute of Information Technology, New Delhi, India.
2. Shubham Choudhury is currently working as Ph.D. in Computational biology from Department of Computational Biology, Indraprastha Institute of Information Technology, New Delhi, India
3. Nisha Bajiya is currently working as Ph.D. in Computational biology from Department of Computational Biology, Indraprastha Institute of Information Technology, New Delhi, India
4. Sumeet Patiyal is currently working as a postdoctoral visiting fellow Cancer Data Science Laboratory, National Cancer Institute, National Institutes of Health, Bethesda, Maryland, USA.
5. Gajendra P. S. Raghava is currently working as Professor and Head of Department of Computational Biology, Indraprastha Institute of Information Technology, New Delhi, India.

## 2. Introduction

In the 1950s, Scholander and his colleagues found fish species that can survive in freezing temperatures, which challenged traditional ideas about living in cold climates [1–3]. In 1969, DeVries and his team linked these adaptations to antifreeze proteins (AFPs) [4]. Since then, AFPs have been discovered in various species, such as insects, fungi, bacteria, and mammals. These proteins help these organisms to survive in extremely cold temperatures either by avoiding freezing or tolerating it [5–10]. AFPs sustain organisms from freezing stress by thermal hysteresis and preventing ice recrystallization [5,11–16]. Due to their unique freeze resistance property, AFPs have a wide range of applications, including food preservation [17–19], medicine [20,21], human cryosurgery, and the production of yoghurt [22–26].

For identifying and characterizing AFPs, numerous experimental techniques have been developed. This includes ammonium precipitation and ice affinity chromatography for purification of AFPs [27–29], Ice-etching [30,31], fluorescence-based ice plane affinity (FIPA) [32], and site-directed mutations [33–35] to determine ice-affinity planes to assess the mechanism of AFPs. These experimental techniques are time-consuming and costly [36]. Simple, similarity-based techniques like BLAST or PSI-BLAST fail to detect distantly related AFPs [23,25,37].

Previously, a number of in silico methods have been used to discriminate AFPs and non-AFPs [38]; a brief description of existing methods is shown in Table 1. In 2011, Kandaswamy and Chou [23] developed AFP-Pred, a pioneered ML-based approach deploying random forest (RF) to classify AFPs. In the same year, Yu and Lu developed an SVM method, iAFP [39], utilising a genetic algorithm (GA) for feature selection. Subsequently, various AFP prediction tools are AFP_PSSM [40], AFP-PseAAC [41], AFP-ensemble [42], TargetFreeze [43], iAFP-Ense [44], CryoProtect [45], afpCOOL [25], and RAFP-Pred [46]. Boosting methods also have been employed by PoGB-pred [47], LightGBM [26], and AFP-LXGB [24]. Recently, sparse representation-based classifiers such as AFP-LSE [16], and AFP-SRC [38,48,49] have also been developed for AFP Prediction.

**Table 1:** List of available methods of AFP prediction.

| Name | Year | Source/Database | Dataset Description |
| --- | --- | --- | --- |
| AFP-Pred* [23] | 2011 | Pfam database | 481 AFPs and 9493 non-AFPs |
| iAFP* [39] | 2011 | PDB,<br>UniProKB | Training: 44 AFPs and 3762 non-AFPs<br>Testing: 369 AFPs |
| AFP_PSSM* [40] | 2012 | AFP-Pred | 481 AFPs and 9493 non-AFPs |
| AFP-PseAAC [41] | 2014 | AFP-Pred | 481 AFPs and 9493 non-AFPs |
| TargetFreeze* [43] | 2015 | AFP-Pred,<br>Pfam database | Training: 300 AFPs and 300 non-AFPs<br>Testing: 181 AFPs and 8293 non-AFPs |
| AFP-ensemble* [42] | 2015 | AFP-Pred | Training: 371 AFPs and 7266 non-AFPs<br>Testing: 93 AFPs and 1817 non-AFPs |
| CryoProtect [45] | 2016 | AFP-Pred | 478 AFPs and 9,139 non-AFPs |
| RAFP-Pred* [46] | 2018 | AFP-Pred,<br>iAFP,<br>UniProKB | Data1: 481 AFPs and 9,493 non-AFPs (AFP-Pred)<br>Data2: 44 AFPs and 3,762 non-AFPs (iAFP)<br>Data3: 369 AFPs (Uniprot) |
| afpCOOL* [25] | 2018 | AFP-Pred,<br>UniProtKB | Data1: 481 AFPs and 9493 non-AFPs<br>Data2: 517 AFP and 517 non-AFPs |
| AFP-CKSAAP [50] | 2019 | AFP-Pred,<br>Pfam | 481 AFPs and 9493 non-AFPs |
| W-GDipc-LRMR-Ri [36] | 2019 | AFP-Pred,<br>MemType-2L | 1. 480 AFPs and 374 no-AFPs<br>2. 7582 MemType-2L dataset |
| AFP-LSE [16] | 2020 | AFP-Pred | 481 AFPs and 9493 non-AFPs |
| Sun et al. [51] | 2020 | AFP-Pred | 481 AFPs and 9493 non-AFPs |
| AFP-CMBPred [52] | 2021 | AFP-Pred, Pfam<br>database | Training: 300 AFPs and 300<br>non-AFPs<br>Testing: 181 AFPs and 8293<br>non-AFPs |
| AFP-LXGB* [24] | 2022 | AFP-Pred, Pfam<br>database | 481 AFPs and 9193 non-AFPs |
| AFP-SPTS* [53] | 2022 | AFP-Pred, Pfam<br>database | 481 AFPs and 9193 non-AFPs |
| AFP-SRC [38] | 2022 | AFP-Pred | 481 AFPs and 9493 non-AFPs |

\*webservice is not working properly or not available.

The major limitations of the existing methods is their dataset, as these methods have been evaluated on unreviewed data. A systematic effort has been made in this study to create a validation dataset of AFPs obtained from Swiss-Prot (reviewed) for evaluation. We implement a robust strategy for AFP prediction by leveraging diverse approaches, including machine learning, sequence similarity, and pattern detection. In addition, ensemble methods have been developed where two or more than two approaches have been integrated to predict AFPs.

## 3. Material and Methods

### 3.1. Dataset compilation and pre-processing

The reliability of a method depends on the quality of the dataset used for training and evaluation. Ideally, both training and testing datasets should be experimentally validated [54,55]. Here, in this study, we have used two datasets, the main and the validation, which were used for model training and external validation, respectively.

#### 3.1.1. Validation Dataset

To create an independent/validation dataset, we extracted AFPs from reviewed sequences of Swiss-Prot [56]. We are left with 81 AFPs length between 16 and 2439 after pre-processing. We have randomly selected the same proportion of negative datasets from Swiss-Prot sequences and left with 73 non-AFPs. Finally, we have 154 validation sequences for model evaluation. We also make sure that the validation data contains only unique sequences.

#### 3.1.2. Main dataset

For the training of our model, we construct the AFPs (positive) dataset by initially obtaining 49352 (unreviewed) AFP sequences from the UniProt database [56]. Additionally, we apply the standard practices followed in the bioinformatics, employing the CD-HIT program [57] to reduce the sequence identity to 40%. After eliminating identical sequences and sequences that contain non-natural amino acids, we were left with 8134 positive sequences with lengths varying from 37 to 12,385. Consequently, the 9493 non-AFPs (negative) were collected from the previously used AFP-Pred. We are left with 9439 non-AFPs after removing the non-natural amino acids. Finally, we have 17573 training sequences for model training and optimization.

### 3.2. Composition Analysis

In this study, we have conducted an assessment of amino acid composition (AAC) for both positive and negative datasets, with a specific focus on AFPs and non-AFPs. Amino acid composition (AAC) serves as the percentage frequency of the 20 different amino acids within a peptide or protein sequence. We have utilized the “Pfeature” [58] module to compute and calculate the AAC composition. To calculate AAC, we applied the following equation:

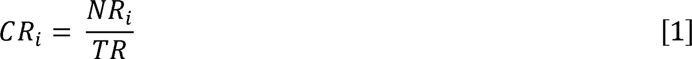

CR_i_ represents the composition of residue i; NR_i_ is the total number of residues of type i; TR stands for the total number of residues in the dataset.

This analysis helps in gaining important insights into the significance of amino acid composition for a better understanding of the quantity and distribution of particular amino acids across various datasets.

### 3.3. Feature generation techniques

Discovering the discriminative features using appropriate techniques is important for designing an effective computational model for classifying the AFPs. In this regard, we have utilized the Pande et al. standalone software “Pfeature” [58] for exploring the salient patterns from primary sequences of AFPs [24].

#### 3.3.1. Composition based features

In this study, we have utilized the composition-based feature module of “Pfeature” to compute the features/descriptors listed in Table 2. This module calculates various features based on the dataset’s composition or proportion of specific elements.

**Table 2:** List of descriptors calculated using the Pfeature.

| Type of Feature | Vector size |
| --- | --- |
| AAC (Amino acid composition) | 20 |
| DPC (Dipeptide composition) | 400 |
| ATC (Atomic composition) | 5 |
| BTC (Bond composition) | 4 |
| PCP (Physico-chemical properties based composition) | 30 |
| RRI (Residue repeat information) | 20 |
| DDOR (Distance distribution of residues) | 20 |
| SER (Shannon entropy for all residues) | 20 |
| SEP (Shannon Entropy at Protein Level) | 1 |
| CTC (Conjoint triad calculation) | 343 |
| PAAC (Pseudo amino acid composition) | 21 |
| APAAC (Amphiphilic pseudo amino acid composition) | 23 |
| QSO (Quasi-sequence order) | 42 |
| SOCN (Sequence order coupling number) | 2 |
| CeTD (Composition enhanced Transition and Distribution) | 189 |
| SPC (Shannon Entropy at Property Level) | 25 |
| Total vector size | 1165 |

#### 3.3.2. Evolutionary based features

The protein’s evolutionary features are known to provide additional crucial information about proteins [59,60]. The evolutionary information of proteins was retrieved from a position-specific scoring matrix (PSSM) profile generated using Position-Specific Iterated BLAST (PSI-BLAST) [61]. In this study, we have used the “pssm_composition” module from the POSSUM package [62] to generate the PSSM-400, a 20 x 20 dimension vector for a protein sequence composition profile that measures the occurrence of 20 amino acids in the sequence. By using the Swiss-Prot, we generated a PSSM matrix for each sequence whose hit has been found in the database. We have created an empty PSSM-400 with zero values for those sequences whose PSSM is not generated due to lack of homologous sequence.

### 3.4. Feature selection techniques

In the majority of datasets, only a small number of features are relevant that contribute to identifying the endpoint. However, a vast number of irrelevant and redundant features remain a significant issue. To overcome this issue, we have applied the feature selection techniques with the aim of identifying the relevant subset of features from the original set. In addition, numerous techniques for feature selection have been developed so far and effectively implemented in various fields [63]. In this study, initially we ranked the features based on the mutual information by implementing an ensemble minimum redundancy–maximum relevance (mRMR) technique [64]. The top-ranked features were most relevant to discriminating AFPs and non-AFPs and complementary to each other [51].

### 3.5. Selection of appropriate ML models

To select an appropriate classifier for the prediction of AFPs, we have used different machine learning algorithms, as its prediction performance will depend not only on the feature representation method but also on the classifier used [24,43]. This study used a different set of classifiers from the scikit-learn [65] library, including k-nearest neighbor (KNN), random forest (RF), logistic regression (LR), gaussian naïve Bayes (GNB), extremely randomized tree (ET), multi-layer Perceptron classifier (MLP) and extreme gradient boosting (XGB).

### 3.6. Cross-Validation Technique

We have adopted the k-fold cross-validation (CV) as a model selection criterion in which the whole dataset is divided into k-folds. In our study, k = 5, then k-1 folds are used for model construction, and the hold-out fold is allocated to model validation [66]. We have divided the complete dataset into 80:20 ratios, where 80% constitutes the training dataset, and 20% constitutes the validation dataset. The model was trained and evaluated based on five-fold cross-validation and the independent validation dataset [51]. Finally, the mean of the performances of five iterations is used as the overall performance [60,67].

### 3.7. Evaluation parameters

We have utilized libraries like Scikit-learn (sklearn), pandas, and NumPy to build machine learning models based on classification. From sklearn, we have imported the following classifiers: RF, ET, GB, MLP, and LR. XGB was imported from XGBoost. Other libraries were installed using the pip packages [68]. In order to evaluate our model’s performance, we used the threshold-dependent metrics of sensitivity, specificity, accuracy, F1-score, kappa, and Mathews correlation coefficient (MCC), and threshold-independent metrics of area under the receiver operating characteristics curve (AUROC) [69]. These threshold-dependent parameters can be calculated as follows:

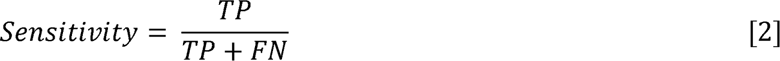

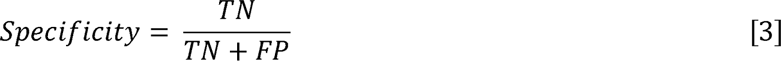

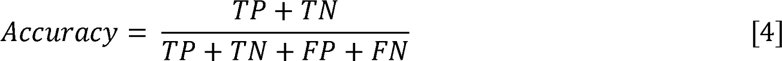

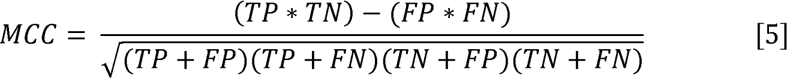

Where, TP stands for true positive; TN stands for true negative; FP stands for false positive; FN stands for false negative

Once the model was evaluated, we chose our top-performing model for further analysis, in which we integrated the evolutionary features with composition-based features and the ML score with the BLAST score and named the hybrid methods

### 3.8. Motif based features

It is crucial to identify conserved motifs in biological sequences in order to discover common shared functions [70]. This study uses the MERCI tool to identify degenerate motifs based on a given classification of amino acids according to their physico-chemical properties [70]. MERCI is employed to select distinct motifs from the positive dataset by analysing input sequences of negative and positive sets. We implemented several MERCI parameters to extract motifs that are exclusive/inclusive to both sets. Exclusive motifs, i.e., not shared among the positive and negative sets, are obtained using the default parameter of the maximal frequency of the negative sequences (fn) as zero. To achieve inclusive motifs, we increased this number to fn = 8. We devised motifs from exclusive and inclusive motifs by determining values such as a) No gap, b) Gap = 1, c) Gap = 2, and d) Class = Koolman-Rohm. Following that, the unique proteins possessing the motifs were identified to estimate the total coverage of motifs in the protein sequence[60].

### 3.9 Webserver Architecture

In this study, we have developed a webserver named “AFProPred” (available at https://webs.iiitd.edu.in/raghava/afpropred/) for the prediction of AFPs and non-AFPs. The front end and back end of webserver is developed using HTML5, CSS, Java, and PHP scripts. It is user-friendly, easily accessible, and compatible with almost all devices, including the desktop, tablet, and mobile phone. In the webserver, we have incorporated 4 major modules such as (i) Predict, (ii) Design, and (iii) Scan.

## 4. Results

In this study, we have divided the result section broadly into the following sub-sections; (i) Compositional analysis of AFPs, (ii) Alignment-based method, (iii) Alignment-free method, (iv) Hybrid approaches, (v) Benchmarking, and (vi) Webserver. A complete workflow of the study is shown in Figure 1, and a description of these subsections can be found below.

**Figure 1:**
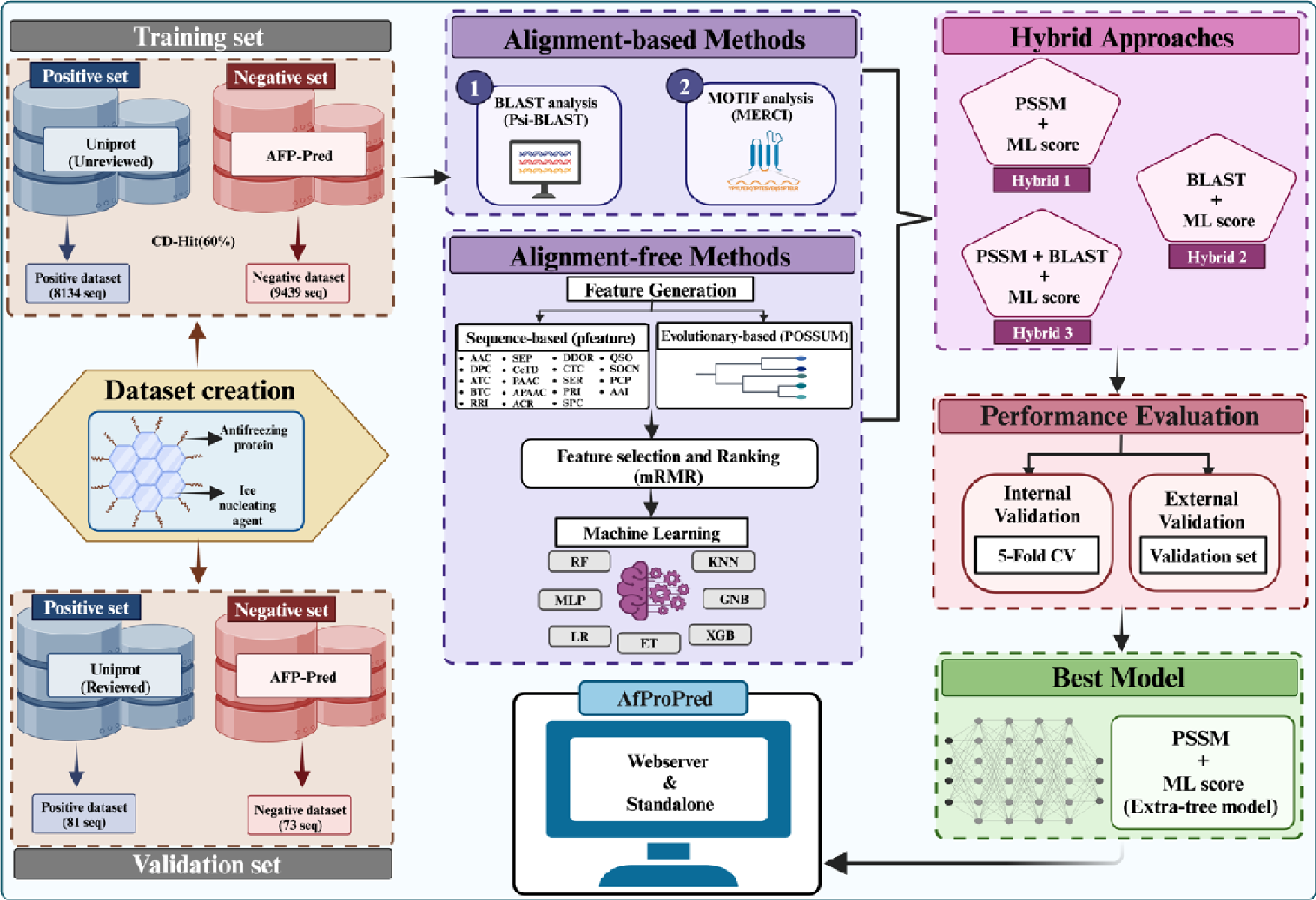
Complete workflow of the study.

### 4.1. Compositional analysis

We have calculated the composition of individual amino acids to assess the prevalence of amino acid residues in each dataset. By analysing the average composition of amino acids in protein sequence, the researcher can identify the potential AFPs. Here, we computed the average residue composition for APFs, non-AFPs, and general proteome in Figure 2, which exhibits that Alanine (A), Isoleucine (I), Valine (V), Threonine (T) are most abundant in AFPs (Positive dataset) while in non-AFPs Leucine (L), Glutamic acid (E) are highly conserved.

**Figure 2:**
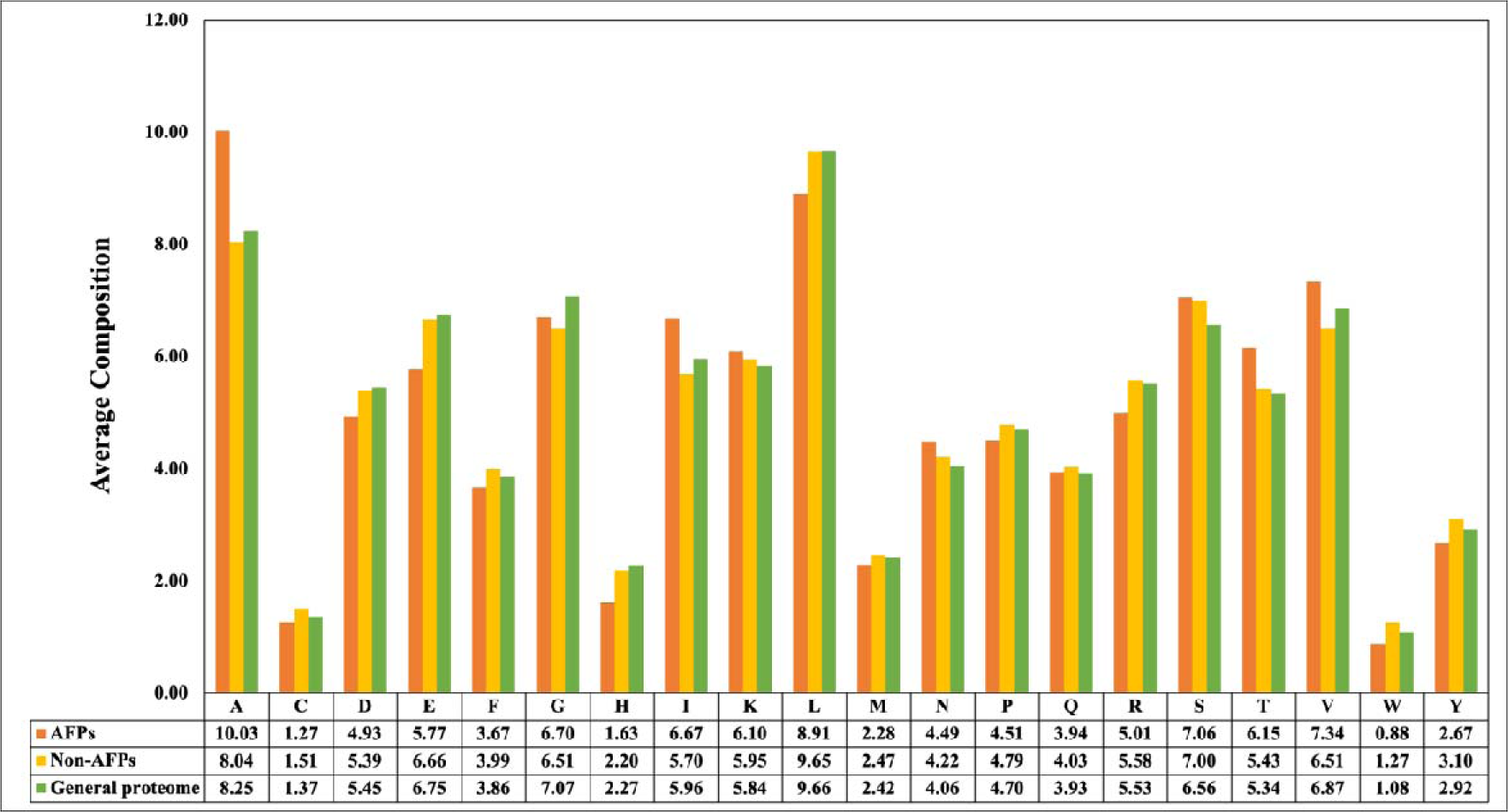
Comparison analysis of the average composition of amino acid residues in AFPs, non-AFPs, and general proteome.

### 4.2. Alignment-based method

#### 4.2.1. Motifs Analysis

Our aim is to identify exclusive and inclusive motifs/patterns present in AFPs, which can be used to search AFPs. We employed the publicly available MERCI software to discover motifs. As MERCI provides number of options, we explore following options or parameters; no gap (default), gap of 1, gap of 2, and Koolman-Rohm (Supplementary Table S1). It’s crucial to remember that MERCI utilizes an fn (maximal frequency in negative sequences) value of zero by default, producing exclusive motifs. We changed the fn value to 8 to widen our analysis and get an inclusive variety of motifs for both negative and positive datasets.

#### 4.2.2. Similarity search approach

A commonly used software package pBLAST (BLAST+ 2.7.1) has been used for performing similarity search where a query sequence is searched against a database [71]. A train dataset was used to build a target database for performing blastp search, where query sequences (sequences in the test set) were searched at different e-values. Based on the top hit, each query sequence has been classified as either AFP or non-AFP. For example, the query sequence is designated as an AFP if the query sequence has the top hit against an AFP. Otherwise, it is assigned as a non-AFP.

### 4.3. Alignment-free method

In this study, various machine learning techniques were applied to construct models for predicting antifreeze proteins [54]. These techniques include Random Forest (RF), Multi-layer Perceptron classifier (MLP), Logistic Regression (LR), XGBoost (XGB), K-nearest neighbor (KNN), Extra Tree (ET), and Gaussian Naive Bayes (GNB). These models have been developed using; i) Compositional feature, ii) PSSM profile, iii) Combined features and iv) Selected features.

#### 4.3.1. Compositional features

We build machine learning based AFP prediction models using different composition-based features generated using the “Pfeature” standalone software (Table 2). Our finding demonstrated that among a spectrum of ML algorithms, the ET, RF, and XGB performed exceptionally. Specifically, the Extra-Tree classifier consistently attained a maximum AUROC score of 0.87 and 0.89 on AAC and APAAC descriptors, respectively, while the performance metrics on other descriptors for the ET classifier are elaborated in Table 3. See Supplementary Table S2 for an in-depth and comprehensive performance of all classifiers.

**Table 3:**
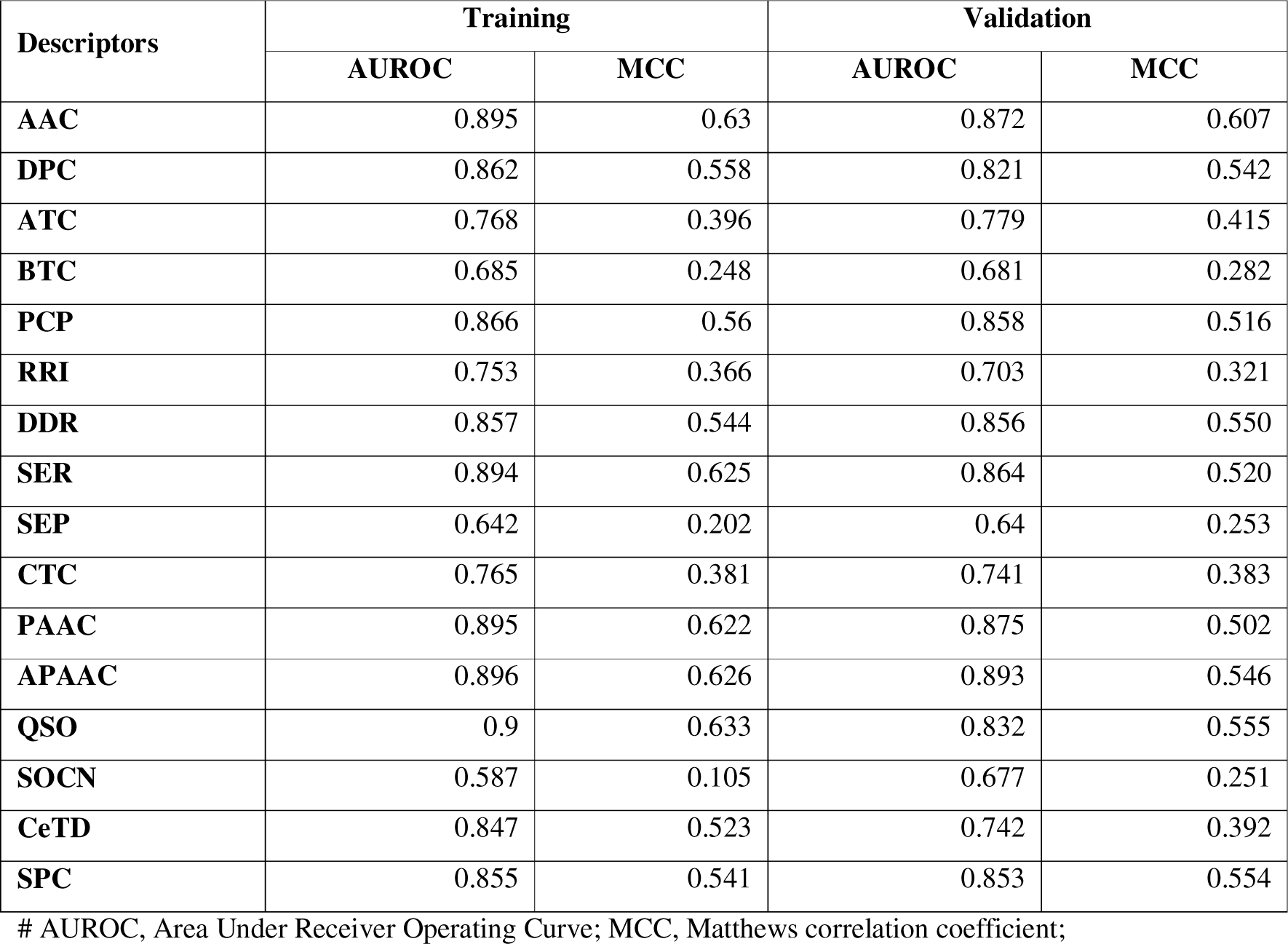
ML performance on different composition-based features.

#### 4.3.2 PSSM profiles

It has been shown in number of studies in the past that evolutionary information based models perform better than single sequence models [72–75]. Thus, this study also explores the potential of evolutionary information in predicting AFPs. We used the POSSUM package [62] to generate evolutionary information for each protein sequence in the form of a PSSM profile. Among all the models generated using the PSSM profile, the ET-based classifier that (Supplementary Table S3) achieved a maximum AUROC of 0.92. To further enhance the predictive performance of our method, we built hybrid methods by combining the PSSM profile with AAC composition features and 150 feature sets, which were calculated using the Pfeature package.

#### 4.3.3. Combined features

Subsequently, we aggregated all compositional features, resulting in a vector of size 1165 for each sequence across the datasets. We then proceeded to develop the prediction models employing the distinct classifier on the combined feature dataset, meticulously fine-tuning the parameters to optimize the AUROC. Our analysis showed that the Extra Tree classifier (ET) and Extreme gradient boosting (XGB) perform almost equally in terms of AUROC, which is 0.86 and 0.87, respectively. Additionally, they achieved MCC scores of 0.54 and 0.63, correspondingly. Table 4 provides a comprehensive overview of performance metrics, encompassing both threshold-dependent and threshold-independent measures, for all the classifiers that were trained and validated across training and validation datasets.

**Table 4:** Performance of different models on the combined features.

| Classifier | Training |  |  |  |  | Validation |  |  |  |  |
| --- | --- | --- | --- | --- | --- | --- | --- | --- | --- | --- |
|  | Sens | Spec | Acc | AUC | MCC | Sens | Spec | Acc | AUC | MCC |
| **ET** | 81.72 | 81.39 | 81.54 | 0.90 | 0.63 | 71.25 | 82.19 | 76.47 | 0.86 | 0.54 |
| **GNB** | 35.28 | 82.74 | 60.78 | 0.67 | 0.21 | 67.50 | 61.64 | 64.71 | 0.67 | 0.29 |
| **KNN** | 58.78 | 73.87 | 66.89 | 0.72 | 0.33 | 38.75 | 69.86 | 53.60 | 0.55 | 0.09 |
| **LR** | 70.85 | 71.99 | 71.46 | 0.78 | 0.43 | 61.25 | 79.45 | 69.94 | 0.82 | 0.41 |
| **MLP** | 69.98 | 70.01 | 69.99 | 0.76 | 0.40 | 56.25 | 90.41 | 72.55 | 0.78 | 0.49 |
| **RF** | 81.72 | 82.10 | 81.92 | 0.90 | 0.64 | 62.50 | 82.19 | 71.90 | 0.84 | 0.45 |
| **XGB** | 82.70 | 82.53 | 82.61 | 0.91 | 0.65 | 82.50 | 80.82 | 81.70 | 0.87 | 0.63 |

#Sens, Sensitivity; Spec, Specificity; Acc, Accuracy; AUROC, Area Under Receiver Operating Curve; MCC, Matthews correlation coefficient; RF, Random Forest; LR, Logistic Regression; XGB, XGBoost; KNNs, k-nearest neighbors; GNB, Gaussian Naive Bayes; ET, Extra-Trees Classification; MLP, Multi-layer Perceptron classifier

#### 4.3.4. Selected features

Previously, it has been shown that number of feature are not relevant they only increase noise in prediction, thus it is important to use only relevant/important features [54,76]. In order to address this challenge, mRMR [77] feature selection techniques that retain the most informative features have been employed, thereby optimizing the feature set for enhanced model performance. In the previous analysis, it was observed that the ET and XGB classifiers perform almost similarly in classifying the AFPs. Therefore, we have opted for the ET-based classifier for further analysis. We employed the mRMR algorithm to rank features and evaluate performance across different feature sets, as detailed in Table 5. It was observed that the ET classifier demonstrated slightly higher performance, achieving an AUROC of 0.90 when utilizing the 150-feature set.

**Table 5:** ET-based performance on different sets of features using the mRMR approach.

| Features | Training |  |  |  |  | Validation |  |  |  |  |
| --- | --- | --- | --- | --- | --- | --- | --- | --- | --- | --- |
|  | Sens | Spec | Acc | AUROC | MCC | Sens | Spec | Acc | AUROC | MCC |
| **100** | 79.24 | 78.81 | 79.01 | 0.88 | 0.58 | 86.25 | 79.45 | 83.01 | 0.89 | 0.66 |
| **150** | 79.41 | 79.09 | 79.24 | 0.88 | 0.58 | 83.75 | 84.93 | 84.31 | 0.90 | 0.69 |
| **200** | 80.22 | 79.30 | 79.73 | 0.88 | 0.59 | 83.75 | 82.19 | 83.01 | 0.89 | 0.66 |
| **400** | 80.44 | 80.91 | 80.69 | 0.89 | 0.61 | 77.50 | 78.08 | 77.78 | 0.86 | 0.56 |
| **800** | 81.13 | 80.63 | 80.86 | 0.89 | 0.62 | 75.00 | 83.56 | 79.09 | 0.87 | 0.59 |

#Sens, Sensitivity; Spec, Specificity; Acc, Accuracy; AUROC, Area Under Receiver Operating Curve; MCC, Matthews correlation coefficient; ET, Extra-Trees Classification

### 4.4. Hybrid approaches

As demonstrated in the preceding sections, we utilized both alignment-based methods (Motif & BLAST) and alignment-free methods (machine learning techniques). Each approach has its own advantages and drawbacks. Alignment-based methods offer high specificity but poor sensitivity. Their performance relies on the similarity or presence of motifs. On the other hand, machine learning-based models or alignment-free methods are more generalized, with performance unaffected by similarity. To capitalize on the strengths of both alignment-free and alignment-based models, we developed ensemble or hybrid methods. Firstly, we developed ML models using combination of PSSM-based feature and AAC. The hybrid PSSM-based ET models perform better than other models, achieving an AUROC of 0.93 with 0.77 MCC, as shown in Table 6. In addition, we developed machine learning techniques using a combination of 150 selected and the PSSM profile. It was observed that RF and ET attain almost similar performance in terms of AUROC of 0.92, which is nearly similar to the hybrid PSSM+AAC. The complete result is provided in Supplementary Table S4. Secondly, we have combined ET-based models developed using AAC with the BLAST score with the aim of enhancing the performance of our ET models. We attained a maximum AUROC of 0.89 on the validation dataset at e-value 1.00E-01. Thirdly, we combined the BLAST-based approach with the AAC+PSSM model and achieved a maximum AUROC of 0.90. In this study, we have tried many combinations (results not shown) to develop hybrid models. We achieved maximum performance AUC of 0.93 on validation dataset using PSSM+AAC feature based on ET-model. Ultimately, we used this model for our study to develop standalone package and web server.

**Table 6:** Performance on the PSSM composition profile combined with AAC composition.

| Hybrid (PSSM + AAC) |  |  |  |  |  |  |  |  |  |  |
| --- | --- | --- | --- | --- | --- | --- | --- | --- | --- | --- |
| Training |  |  |  |  |  | Validation |  |  |  |  |
| Classifier | Sens | Spec | Acc | AUR<br>OC | MCC | Sens | Spec | Acc | AUR<br>OC | MCC |
| **ET_new** | 82.91 | 83.66 | 83.32 | 0.92 | 0.67 | 85.00 | 91.78 | 88.24 | 0.93 | 0.77 |
| **GNB** | 51.48 | 78.79 | 66.15 | 0.73 | 0.32 | 46.25 | 91.78 | 67.97 | 0.85 | 0.42 |
| **KNN** | 82.35 | 82.07 | 82.20 | 0.90 | 0.64 | 68.75 | 87.67 | 77.78 | 0.89 | 0.57 |
| **LR** | 80.16 | 80.06 | 80.11 | 0.87 | 0.60 | 47.50 | 79.45 | 62.75 | 0.79 | 0.28 |
| **MLP** | 85.52 | 85.68 | 85.60 | 0.93 | 0.71 | 70.00 | 73.97 | 71.90 | 0.80 | 0.44 |
| **RF** | 83.38 | 83.19 | 83.28 | 0.91 | 0.67 | 76.25 | 87.67 | 81.70 | 0.91 | 0.64 |
| **XGB** | 84.13 | 84.37 | 84.26 | 0.92 | 0.68 | 73.75 | 78.08 | 75.82 | 0.86 | 0.52 |

#Sens, Sensitivity; Spec, Specificity; Acc, Accuracy; AUROC, Area Under Receiver Operating Curve; MCC, Matthews correlation coefficient; RF, Random Forest; LR, Logistic Regression; XGB, XGBoost; KNNs, k-nearest neighbors; GNB, Gaussian Naive Bayes; ET, Extra-Trees Classification; MLP, Multi-layer Perceptron classifier

### 4.5. Benchmarking

In order to assess the significance of a newly developed method, it is important to compare it performance with available methods on same dataset [78]. Thus, we evaluate existing methods on our validation dataset. Unfortunately, all existing methods are not available, thus, we are able to predict performance of those methods that fully functional and available to public. Our findings show that existing methods (CryoProtect, iBLP) don’t perform better on the validation dataset than our method (AFPpropred) as shown in Table 7.

**Table 7:** Comparison of available methods.

| Name | Year | Spec | Sens | Acc | AUROC | MCC |
| --- | --- | --- | --- | --- | --- | --- |
| Cryoprotect | 2017 | 42.50 | 79.45 | 60.13 | 0.61 | 0.23 |
| iBLP | 2021 | 13.92 | 83.56 | 47.36 | 0.48 | -0.03 |
| AFProPred | - | 94.52 | 86.25 | 90.20 | 0.93 | 0.77 |

#Sens, Sensitivity; Spec, Specificity; Acc, Accuracy; AUROC, Area Under Receiver Operating Curve; MCC, Matthews correlation coefficient;

### 4.6. Webserver Implementation

In order to provide a more user-friendly webserver for the scientific community, we developed AFProPred, which can be accessed at https://webs.iiitd.edu.in/raghava/afpropred/. This platform is designed for predicting AFPs using our top-performing model. AFProPred offers several modules, namely Prediction, Design, Protein Scan, and Motif Scan. The Prediction module effectively distinguishes between AFPs and non-AFPs, allowing users to submit protein sequences in FASTA format for prediction. The Design module enables users to generate all possible analogs of input sequence, and predict AFPs among analogs. Protein Scan module assists in identifying AFP regions within a given protein sequence. This platform is developed using responsive template to browse on wide range of devices including, smart phones, laptops and desktops. Additionally, we have created a Python-based standalone package named "AFProPred" to facilitate users in predicting AFP proteins at genome scale. This package is available for download via the "download module" on the web server at https://webs.iiitd.edu.in/raghava/afpropred/standalone.html.

## 5. Discussion

Numerous methods have been developed in the past to improve the accuracy of prediction of AFPs (Table 1). Most of the existing studies used AFP-Pred dataset, which contain 481 AFPs and 9493 non-AFP. AFPs were mainly extracted from Pfam database in few studies (like as iAFP) extracted AFPs from PDB. One of the major limitation of existing studies is their dataset, none of the existing study used experimentally validated AFPs or well annotated AFPs available in Swiss-prot which contain reviewed proteins. Thus validity of these studies is questionable because garbage in results to garbage out. Thus, developing a method based on annotated or experimentally supported data is critical. In this study, we made a systematic attempt to create a dataset of well annotated AFPs. We queried Swiss-prot for extracting annotated AFP and got only 81 AFPs, which is not sufficient for training, testing and validating our prediction models. Thus, we used these 81 AFPs only for validating our models and existing models. Similarly, we obtained 73 non-AFPs from Swiss-prot using keyword “not anti-freezing proteins”. Finally, we build an independent dataset that contain well annotated 81 AFPs and 73 non-AFPs. This independent dataset is not used for training or optimization of hyper parameters of machine learning techniques.

Our primary analysis indicated, common evolutionary relationships among antifreeze proteins where certain residues are conserved in AFPs (e.g., Ala, Ile, Val, and Thr). The amino acid Thr increase the activity of AFPs by the addition of hydrogen bonds on their surface area [45]. Firstly, we utilized similarity based standard technique BLAST to identify AFPs. In this study, BLAST performs fails to discriminate AFPs and non-AFPs due poor similarity among AFPs. In order to overcome this challenge, we used generalized techniques such as machine learning techniques for predicting AFPs. These techniques are also called alignment-free techniques as they are not based on alignment. Most of the machine learning techniques need fixed length vectors whereas proteins have variable length, thus we compute composition based features [16]. Here, we tried wide range of machine learning techniques and achieved maximum AUC 0.90 on independent dataset. It well known fact that evolutionary information is important for predicting function of proteins [79]. Thus in this study also we developed models using evolutionary information which is obtained from PSSM profile generated using PSI-BLAST. Performance of our model also improved from AUC 0.90 to 0.93 when PSSM is used instead of protein composition. Finally, we compare performance of our method with existing dataset. In order to provide fair evaluation, we evaluate performance of our model as well as existing methods on independent dataset of 81 AFPs and 73-non AFPs. As shown in result section, our method perform better than existing methods.

## 6. Conclusion

To uncover the relationship between proteins and ice-crystals as well as, more generally, the adaptation of organisms to their environments depends on the ability to understand the evolution of AFPs [51]. Our findings revealed that the conservation of several essential amino acids showed opposite tendencies in AFPs and non-AFPs. This suggests that there has been a significant selection pressure related to these amino acids leading to the differentiation between AFPs and non-AFPs regarding their ice-binding capacities. Therefore, our hybrid approaches, AAC combined with PSSM profiles, performed significantly and outperformed the state-of-the-art tools; hence, they are an effective and beneficial tool for identifying new antifreeze proteins.

## Abbreviations

AAC: Amino acid composition
Ala: Alanine
AUROC: Area Under the Receiver Operating Characteristic
BLAST: Basic Local Alignment Search Tool
ET: Extra Tree classifier
FS: Feature selection
GNB: Gaussian Naive Bayes
Ile: Isoleucine
KNN: k-Nearest Neighbor
LR: Logistic Regression
MCC: Matthew’s correlation coefficient
ML: Machine learning
MLP: Multi-layer Perceptron classifier
PSI-BLAST: Position-Specific Iterated BLAST
PSSM: Position-Specific Scoring Matrix
RF: Random Forest
Thr: Threonine
Val: Valine
XGB: eXtreme Gradient Boosting

## Authors’ contributions

GPSR, collected the dataset. NK and SP processed the dataset. NK, SP, and GPSR implemented the algorithms and developed the prediction models. NK, NB, and GPSR analysed the results. SC created the front-end, back-end of the webserver and standalone of the method. NK, NB, SP, SC, and GPSR penned the manuscript. GPSR conceived and coordinated the project. All authors have read and approved the final manuscript.

## Acknowledgments

Authors are thankful to the University Grants Commission (UGC), Department of Bio-Technology (DBT), DBT-RA program in Biotechnology and Life Sciences, Department of Science and Technology (DST-INSPIRE), and Council of Scientific & Industrial Research (CSIR) for fellowships and financial support, and the Department of Computational Biology, IIITD New Delhi for infrastructure and facilities. We would like to acknowledge that Figures were created using BioRender.com.

## Data Availability Statement

All the datasets used in this study are available at the “AFProPred” web server, https://webs.iiitd.edu.in/raghava/afpropred/dataset.html.

